# Fluorescent human RPA to track assembly dynamics on DNA

**DOI:** 10.1101/2023.11.23.568455

**Authors:** Vikas Kaushik, Rahul Chadda, Sahiti Kuppa, Nilisha Pokhrel, Abhinav Vayyeti, Scott Grady, Chris Arnatt, Edwin Antony

**Affiliations:** Department of Biochemistry and Molecular Biology, St. Louis University School of Medicine, St. Louis, MO 63104; Department of Biological Sciences, Marquette University, Milwaukee, WI 53233; Department of Chemistry, St. Louis University, St. Louis, MO 63103

**Keywords:** Human Replication Protein A (RPA), 4-azido-*L*-phenylalanine (4AZP), site-specific labeling, RPA Interacting proteins (RIPs), ssDNA, click chemistry, non-canonical amino acids, genetic code expansion

## Abstract

DNA metabolic processes including replication, repair, recombination, and telomere maintenance occur on single-stranded DNA (ssDNA). In each of these complex processes, dozens of proteins function together on the ssDNA template. However, when double-stranded DNA is unwound, the transiently open ssDNA is protected and coated by the high affinity heterotrimeric ssDNA binding Replication Protein A (RPA). Almost all downstream DNA processes must first remodel/remove RPA or function alongside to access the ssDNA occluded under RPA. Formation of RPA-ssDNA complexes trigger the DNA damage checkpoint response and is a key step in activating most DNA repair and recombination pathways. Thus, in addition to protecting the exposed ssDNA, RPA functions as a gatekeeper to define functional specificity in DNA maintenance and genomic integrity. RPA achieves functional dexterity through a multi-domain architecture utilizing several DNA binding and protein-interaction domains connected by flexible linkers. This flexible and modular architecture enables RPA to adopt a myriad of configurations tailored for specific DNA metabolic roles. To experimentally capture the dynamics of the domains of RPA upon binding to ssDNA and interacting proteins we here describe the generation of active site-specific fluorescent versions of human RPA (RPA) using 4-azido-*L-*phenylalanine (4AZP) incorporation and click chemistry. This approach can also be applied to site-specific modifications of other multi-domain proteins. Fluorescence-enhancement through non-canonical amino acids (FEncAA) and Förster Resonance Energy Transfer (FRET) assays for measuring dynamics of RPA on DNA are also described.

**Highlights:**

- RPA is an essential protein for most DNA metabolic processes including replication, repair, and recombination.
- RPA is a ssDNA binding protein made of six domains situated across RPA70, RPA32 and RPA14 subunits. Four high affinity DNA binding domains engage the DNA.
- Site-specific fluorescent probes were incorporated into two domains of RPA and report on ssDNA binding dynamics.
- Bulk-level kinetic and single-molecule assays are described to monitor the binding and remodeling of individual RPA domains on ssDNA.

## Introduction

RPA is an essential single-stranded DNA (ssDNA) binding protein required for most aspects of DNA metabolism including replication, repair, recombination, and telomere maintenance^1-4^. Almost all DNA repair processes required to maintain genomic integrity are coordinated by ssDNA intermediates coated by RPA. The myriad functions of RPA are orchestrated through four key steps^1-4^: i) RPA binds to ssDNA with high affinity (K_D_ <10^−10^M) and protects it from degradation by exo- and endonucleases. ii) Formation of RPA-ssDNA complexes triggers the DNA damage checkpoint response. iii) RPA physically interacts with over three dozen DNA processing enzymes and recruits most of them onto the site of DNA damage. iv) Finally, RPA hands-off the DNA to these enzymes and correctly positions them to facilitate/regulate their catalytic activity. While the absolute requirement for RPA in DNA metabolism and maintenance is well noted, precisely how RPA contributes to such mechanisms is poorly understood.

RPA achieves functional versatility using structural modularity and flexibility (**Figure 1**). RPA70, RPA32 and RPA14 subunits make up the RPA heterotrimer^5^. RPA-DNA and RPA-protein interactions are coordinated through six oligonucleotide/oligosaccharide binding (OB) folds labeled A-F^6-9^, however only four of the OB-folds are predominantly involved in direct DNA interactions^8^. We term these *DNA binding domains* (DBDs). DBDs-A, B and C reside in RPA70 and are connected by flexible linkers. DBD-D resides in RPA32. RPA14 is part of the trimerization core (Tri-C) and holds all three subunits together. There are two *protein interaction domains* (PIDs) which bind to over three dozen enzymes involved in DNA metabolism^3,5,10,11^. One PID is situated at the N-terminus of RPA70 (OB-F or PID^70N^) and is connected by a long flexible linker to DBD-A. A second winged helix (wh) motif containing PID is situated at the C-terminus of RPA32 (wh or PID^32C^) and is connected by a shorter disordered linker. A 40-amino acid region located at the N-terminus of RPA32 is unstructured and is extensively phosphorylated by multiple kinases^4,12^. Because of the flexible linkers the DBDs and PIDs RPA can adopt a multiplex of structural *configurations* (defined as the relative position of one domain with respect to others). This feature has been hypothesized to be the functional basis for how RPA serves multiple roles in the cell^11,13-16^. Differential configurations of the DBDs/PIDs and/or the number of DBDs bound on ssDNA would lead to transient open pockets of ssDNA, onto which RPA-interacting proteins (RIPs) can be loaded^2,11^.

**Figure 1.**
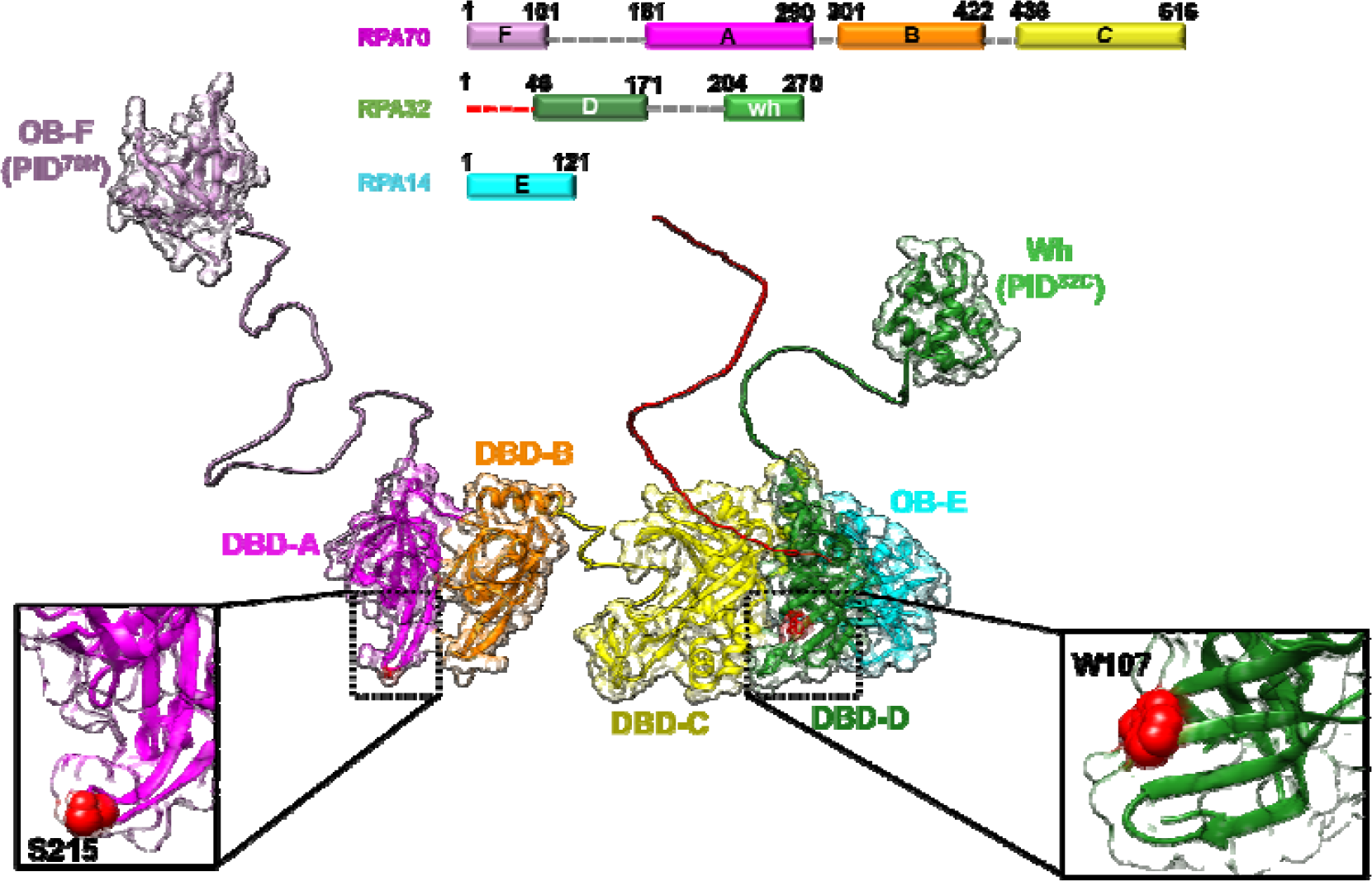
Structural features of hetero-trimeric human RPA. RPA70, RPA32, and RPA14 subunits and the individually colored oligonucleotide/oligosaccharide binding (OB) domains are shown. The disordered linkers between the various domains are also depicted. Position of S215 in DBD-A (RPA70) and W107 in DBD-D (RPA32) are shown as inserts and are positions used for incorporation of 4-azido-*L*-phenylalanine and attachment of fluorescent dyes through click chemistry. The structure was assembled using AlphaFold2 predicted models AF-P27694-F1 (RPA70), AF-P15927-F1 (RPA32), and AF-P35244-F1(RPA14) in combination with high resolution structures of isolated RPA70 (181-422; PDB:1JMC), RPA32 and RPA14 (PDB:2PI2) domains.

A persistent problem in deciphering how certain RPA configurations drive specific cellular functions arises from experimental difficulties. Since there are multiple moderate-affinity ssDNA binding domains, RPA is always macroscopically bound to ssDNA even when mutations are introduced within a particular DBD^2,17,18^. Thus, changes in overall K_D_ are typically not observed. To circumvent this problem, the DNA binding properties of isolated domains were traditionally used for structure-function investigations^2,11^. DBDs A and B were purified separately, or together, and their ssDNA binding constants were determined^19^: K_D_ = 2 μM (DBD-A) ^13^, 16.8 μM (DBD-B) ^13^, 52 nM (DBDs A+B) ^5,20^. In comparison, full length RPA binds to ssDNA with even higher affinity (K_D_ <0.1 nM) ^19,21,22^. Isolated trimerization core, made up of DBD-C (RPA70), DBD-D (RPA32), and RPA14, is thought to have weaker ssDNA binding affinity (K_D_ ∼5 μM)^20,23-26^. Unfortunately, effects of allostery and inter-domain interactions on ssDNA binding are lost in such studies. A second approach has utilized mutagenesis of select Trp residues within individual DBD to create Aro-mutants^27^. These Trp residues base-stack with ssDNA and contribute to binding but do come with the caveat of perturbing other key interactions. In addition, several electrostatic interactions also contribute to DNA binding. Based on such studies, existing models for RPA assume that the DBD-A-B pair form a high affinity ssDNA binding module, whereas the trimerization core act as a low affinity module. However, structural studies of RPA show that the DBD-A-B module binds to 9 nucleotides whereas the trimerization core binds to 14 nucleotides (nts)^8,28^. Thus, the K_D_ estimated from studies of isolated DBDs-ssDNA interactions are at odds with the structural evidence.

To uncover the contributions of the individual DBDs to the overall DNA binding mechanism, a unique signal/signature from within each DBD is required upon DNA binding. We site-specifically positioned the non-canonical amino acid 4-azido-*l-*phenylalanine (4AZP) at either DBD-A or DBD-D and tethered fluorophores using strain promoted cycloaddition click chemistry. We originally developed the methodology for *S. cerevisiae* RPA and uncovering the binding dynamics of these individual DBDs on ssDNA from within the context of the full-length protein^17,18,28^. Here we describe the development of this methodology for human RPA and showcase the power of the approach through a few ensemble and single molecule experimental examples.

### Approach

Protein-induced fluorescence enhancement (PIFE) produces a change in fluorescence quantum yield upon proximal substrate binding when a fluorophore is positioned either on the protein of interest or on the substrate (e.g., DNA or RNA)^29-31^. Positioning the fluorophore on substrates such as nucleic acids is relatively straightforward as the modifications can be site-specifically introduced during oligonucleotide synthesis. For proteins, the challenge arises from the nuances required to incorporate and position the fluorophore at a single position. For RPA, three objectives must be met: a) A single fluorophore must be inserted within one DBD. b) A change in fluorescence must be generated when the DBD binds to DNA. C) There should be no effects on DNA binding or protein-protein interactions. Cys-based maleimide chemistry is not suitable for site-specific labeling of large proteins like RPA. There are 1007 amino acids and 15 Cys residues scattered across the three subunits. Thus, generating a single-Cys version without loss in RPA activity is a daunting task. Screening for an accessible Cys and obtaining reproducible uniform labeling are impossible when multiple Cys residues are present. Thus, we utilized non-canonical amino acids and engineered 4-azido-*L*-phenylalanine (4AZP) at single positions in two specific DBDs ^18,32-36^. Next, using strain promoted alkyne-azide cycloaddition (click chemistry) ^37^, we covalently conjugated fluorophores such as MB543, Cy3, or Cy5. Optimal positioning of MB543 at position S215 in RPA70 (DBD-A) or W107 in RPA32 (DBD-D) produced PIFE changes upon ssDNA binding. More importantly, extensive functional characterization revealed no loss in DNA binding or protein-interaction activities. We term this approach <u>F</u>luorescence <u>E</u>nhancement through <u>N</u>on-<u>C</u>anonical <u>A</u>mino <u>A</u>cids (FEncAA)^38^. A detailed procedure to generate fluorescent human RPA using non-canonical amino acids and click chemistry is elaborated below.

## Materials and Methods

### Plasmids for protein overexpression and 4AZP incorporation

A plasmid expressing all three subunits of RPA (p11d-RPA) was a kind gift from Dr. Marc Wold (University of Iowa). This plasmid was modified using Q5-site directed mutagenesis to generate two separate plasmids for generation of fluorescent RPA modified at either DBD-A (RPA70) or DBD-D (RPA32). First, all amber STOP codons (TAG) were changed to ochre stop codons (TAA) in p11d-RPA. RPA-S215TAG-p11d was generated by engineering a C-terminal 6X-polyhistidine tag in RPA70 along with a TAG substitution at the position coding for Ser215 in RPA70. RPA-W107TAG-p11d was generated by engineering a C-terminal 6X-polyhistidine tag in RPA32 along with a TAG substitution at the position coding for Trp107 in RPA32. The TAG stop codons mark the site for incorporation of *p*-azido-L-phenylalanine (4AZP). The pDule2-pCNF plasmid carrying the tRNA and tRNA Synthetase specific for 4AZP incorporation at the TAG specific positions was a kind gift from Dr. Ryan Mehl (Oregon State University).

### Synthesis of 4AZP

We previously detailed a procedure for the synthesis of 4AZP from Fmoc-4-amino-phenylalanine ^38^. We here describe a modified procedure with minor changes but results in more consistent 4AZP yields. 25 g (62.1 mmol) of Fmoc-4-amino-phenylalanine (Angene International Ltd., China) was added to 2 L D.I. water in a 4 L Erlenmeyer flask with a stir bar. The flask was placed in an ice bath and the suspension was stirred while concentrated HCl (37% v/v, 30 mL) and NaNO_2_ (5.0 g, 73 mmol dissolved in 30 mL of water) were added sequentially over a period of 20 min. The resulting off-white suspension was stirred at 4 °C for 3 h, followed by addition of sodium azide (5.05 g, 77.7 mmol, dissolved in 20 mL D.I. water), dropwise over a period of 20 min. This reaction was allowed to proceed overnight in the dark. Next, 2 L dichloromethane (in 500 mL batches) was added to the overnight suspension with stirring. Aqueous and organic phases were separated using a 2 L separatory funnel. The organic layer containing azide 1 was concentrated using a rotary evaporator to a final volume of 1 L. Next 120 mL of piperidine (1.2 mol) was added to the final suspension for an overnight reaction at room temperature (with stirring). Next, 1 L of 1N NaOH (pellets dissolved in water) was added to the DCM suspension and stirred for 30 min. The top aqueous layer was separated from DCM using 2 L separatory funnel and washed with 3 batches (250 mL each) of DCM to remove residual piperidine. The pH of the alkaline aqueous layer was adjusted to 7.0 using 10 M aqueous HCl. It is imperative to maintain the pH during this step and thus please add HCl in small increments as the product will form at pH 7.0. Periodically monitor changes in pH and readjust as needed. The resulting orange suspension was concentrated to 500 mL using a rotary evaporator at 60 °C. H_2_O-HCl mixture (pH 0.8) heated at 70 °C was added to the concentrated suspension (heated at 70 °C) until majority of the product went into solution. Hot filtration was performed for crystallization of the product and the filtered suspension was allowed to cool at room temperature. This step is best performed using a side-arm flask under vacuum. Light brown 4AZP crystals will be visible immediately after filtration. The crystallization mixture was covered with tinfoil and kept at 4 °C. Crystals, formed overnight, were filtered under vacuum while still cold. Filtered crystals were freeze-dried, distributed in glass vials, and stored at 4 °C for further use. A lyophilizer was used for drying the 4AZP.

### Protein Expression and Purification

600 ng of each plasmid (RPA-W107TAG-p11d or RPA-S215TAG-p11d and pDule2-pCNF) were co-transformed using heat shock competent BL21(DE3)pLysS *E. coli* cells. Please note that selection of cell line is imperative for optimum 4AZP incorporation and final yield of protein. BL21(DE3)pLysS cells work best for human RPA. If the methodology is used for other proteins, we recommend screening multiple cell lines for expression and yield. Successful transformants were selected on LB agar plates containing ampicillin and spectinomycin and grown for 12-16 h at 37 °C until colonies appeared. For optimal RPA protein overexpression and 4AZP incorporation we recommend using freshly transformed cells as opposed to glycerol stocks. Ampicillin and spectinomycin were used at 100 and 50 μg/mL final concentration, respectively.

A single colony was added to 1 L minimal media. The media was prepared by adding the following components to a 2.8 L baffled Fernbach flask. 814.6 mL of D.I. water, 50 mL of 10% glycerol, 1.2 mL of 40% glucose, 20 mL of 50X M salts (**Table 1**), 2 mL of 1 M MgSO_4_, 50 mL of 5% aspartate pH 7.5, 20 mL of 4 mg/mL leucine pH 7.5, 40 mL of the 25X amino acids mix (**Table 2**), 200 μL of 5000X trace metal stock solution, 1 mL of 100 mg/mL ampicillin, and 1 mL of 50 mg/mL spectinomycin. Please note that individual metal stocks can be prepared in 30 mL batches (**Table 3**). Autoclave all solutions, except for CuCl_2_ and FeCl_3_ which tend to precipitate when autoclaved and thus need to be filtered using a 0.22 μm syringe filter. The metal stock solutions can be stored at room temperature. Using the metal stocks, prepare a 50 mL 5000X trace metal stock solution (**Table 4**). Since RPA overproduction is toxic to *E. coli* cells, after adding the colony, we recommend incubating the media for 16-20 h without shaking at 37 °C. This step significantly improves overall protein yield. After overnight incubation, begin shaking at 250 rpm until OD_600_ reaches 1.8-2.0, induce with 0.4 mM IPTG and add freshly prepared 1 mM 4AZP (206 mg of 4AZP crystals is dissolved in 6 ml of water and then added to each 1L media). Cells were then allowed to grow at 37 °C for an additional 3 h with shaking at 250 rpm. After 3 h of induction, cells were harvested by centrifugation at 3,470 xg for 30 min. Supernatant was decanted and the cell-pellet was resuspended in 25 mL of cell resuspension buffer (30 mM HEPES, pH 7.8, 300 mM KCl, 0.02% Tween-20, 10% glycerol (V/V), 3X protease inhibitor cocktail (PIC). This slurry can be flash frozen and stored at -80 °C for 4-6 weeks.

**Table 1.** 50X M Salt solution (autoclave and store at room temperature)

| 50X M salts components | Weight (g) per 500 mL | Final Concentration (M) |
| --- | --- | --- |
| $\text{Na}_2\text{HPO}_4$ | 88.73 g | 1.25 M |
| $\text{KH}_2\text{PO}_4$ | 85.1 g | 1.25 M |
| $\text{NH}_4\text{Cl}$ | 66.9 g | 2.5 M |
| $\text{Na}_2\text{SO}_4$ | 17.8 g | 0.25 M |

**Table 2.**
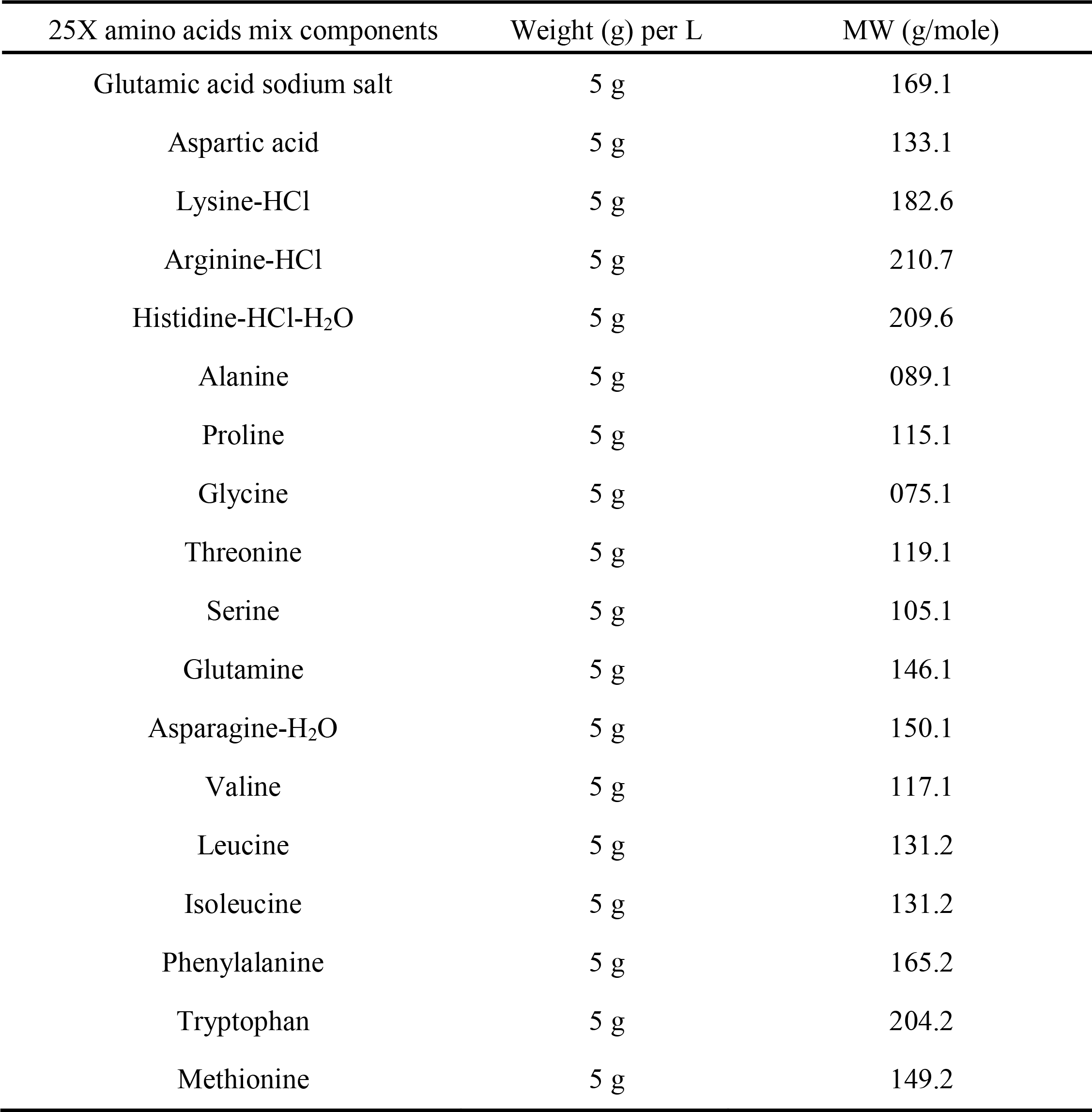
25X Amino acids mix (filter via 0.22 μm store at 4 °C)

**Table 3.** Individual Trace Metal Solutions.

| Metal | Amount for 30 mL solution |
| --- | --- |
| CaCl<sub>2</sub>.2H<sub>2</sub>O | 8.82 g |
| MnCl<sub>2</sub>.4H<sub>2</sub>O | 5.93 g |
| ZnSO<sub>4</sub>.7H<sub>2</sub>O | 8.62 g |
| CoCl<sub>2</sub>.6H<sub>2</sub>O | 1.32 g |
| CuCl<sub>2</sub> | 807 mg |
| NiCl<sub>2</sub> | 777 mg |
| Na<sub>2</sub>MoO<sub>4</sub> | 1.45 g |
| Na<sub>2</sub>SeO<sub>3</sub> | 1.03 g |
| H<sub>3</sub>BO<sub>3</sub> | 371 mg |
| FeCl<sub>3</sub> | 486 mg |

**Table 4.**
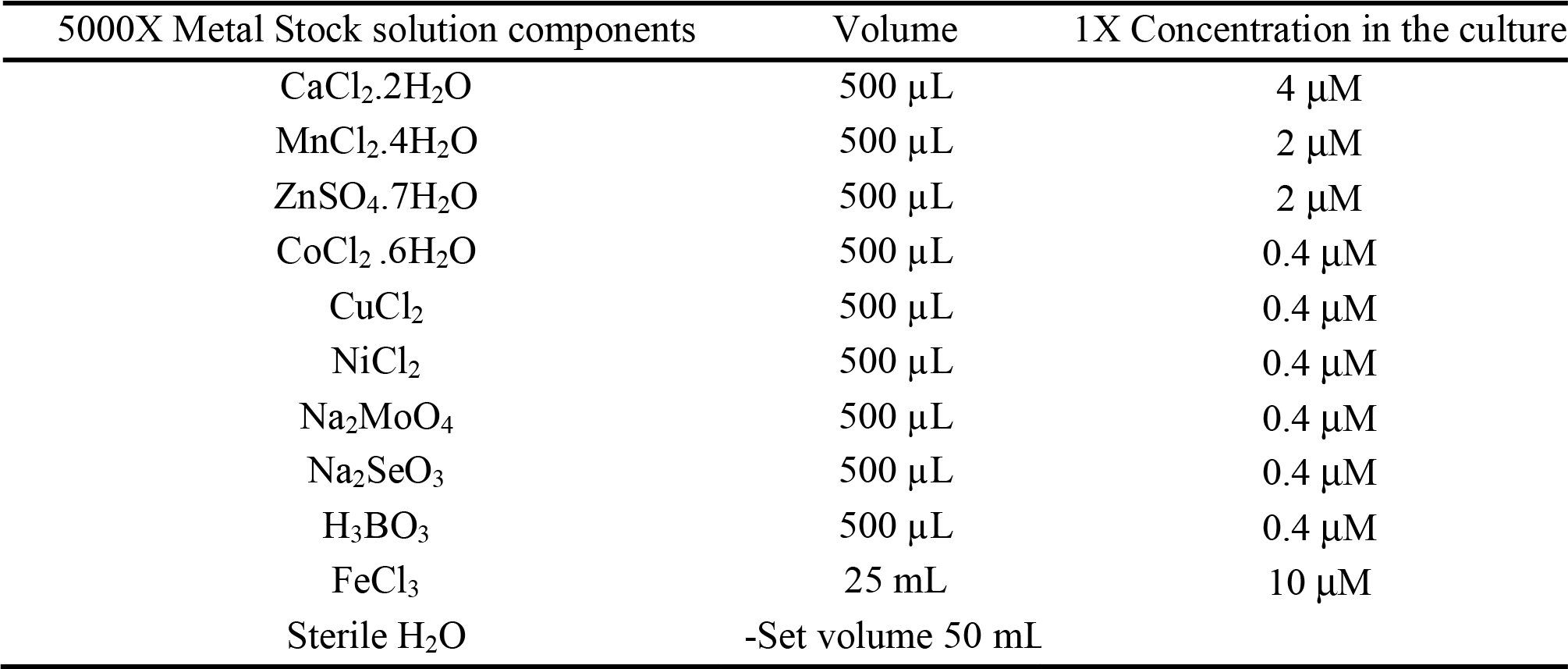
5000X Trace Metal Stock solution.

Cell lysis was performed using 0.4 mg/mL lysozyme in the presence of 1 mM phenylmethylsulfonyl fluoride (PMSF) and incubated at 4 °C for 30 min with gentle stirring. The lysate was sonicated at 50% amplitude for a total of 2 min with 50% duty cycle with each cycle lasting for a minute. After centrifugation for 1 h at 37,157 xg, the clarified supernatant containing RPA^*4AZP*^ was applied to a 3 mL pre-equilibrated Ni^2+^-NTA (Gold Biotechnology Inc., USA) gravity column. The resin was washed sequentially to remove nonspecifically bound proteins with 100 mL each of Ni^2+^ -NTA wash buffer 1 (30 mM HEPES, pH 7.8, 300 mM KCl, 0.02% Tween-20, 10% glycerol (V/V), 10 mM imidazole, 1.5X PIC), Ni^2+^ -NTA wash buffer 2 (30 mM HEPES, pH 7.8, 2 M KCl, 0.02% Tween-20, 10% glycerol (V/V), 10 mM imidazole, 1.5X PIC), and Ni^2+^ -NTA wash buffer 3 (30 mM HEPES, pH 7.8, 50 mM KCl, 0.02% Tween-20, 10% glycerol (V/V), 10 mM imidazole, 1.5X PIC). Bound protein was eluted with 20 mL of Ni^2+^ -NTA elution buffer (30 mM HEPES, pH 7.8, 300 mM KCl, 0.02% Tween-20, 10% glycerol (V/V), 400 mM imidazole, 1.5X PIC) and eluates were assessed for RPA content on a 12% SDS-PAGE gel (**Figure 2**). Fractions containing RPA^*4AZP*^ were pooled and the conductivity adjusted with H_0_ buffer (30 mM HEPES, pH 7.8, 0.02% Tween-20, 10% glycerol (V/V), 0.25 mM EDTA, pH 8.0, 1.5X PIC) to match the conductivity of H_100_ buffer (30 mM HEPES, pH 7.8, 100 mM KCl, 0.02% Tween-20, 10% glycerol (V/V), 0.25 mM EDTA, pH 8.0, 1.5X PIC). The conductivity adjusted pool was fractionated further over a 5 mL HiTrap Heparin column equilibrated with H_100_ buffer. The nonspecifically bound proteins were washed off with 100 mL of H_100_ buffer, followed by elution with 100 mL gradient of H_100_ to H_1.5M_ buffer (30 mM HEPES, pH 7.8, 1.5 M KCl, 0.02% Tween-20, 10% glycerol (V/V), 0.25 mM EDTA, pH 8.0, 1.5X PIC). Fractions containing RPA^*4AZP*^ (**Figure 2 C & D**) were identified by analyzing the fractions on a 12% SDS-PAGE gel, pooled, and concentrated. As a final purification step, the sample was fractionated over a HiLoad 16/ 600 Superdex 200 pg size exclusion column using RPA Storage buffer (30 mM HEPES, pH 7.8, 200 mM KCl, 0.01% Tween-20, 10% glycerol (V/V), 0.25 mM EDTA, pH 8.0). All FPLC buffers were filtered and degassed. Fractions containing RPA^*4AZP*^ were confirmed on a 12% SDS-PAGE gel, pooled, and concentrated (**Figure 2 E & F**). RPA^*4AZP*^ concentration was determined using extinction coefficient 87,410 M^-1^cm^-1^ at 280 nm. The protein was flash frozen as single use aliquots (10 μM; 0.5 mL) and stored at -80 °C.

**Figure 2:**
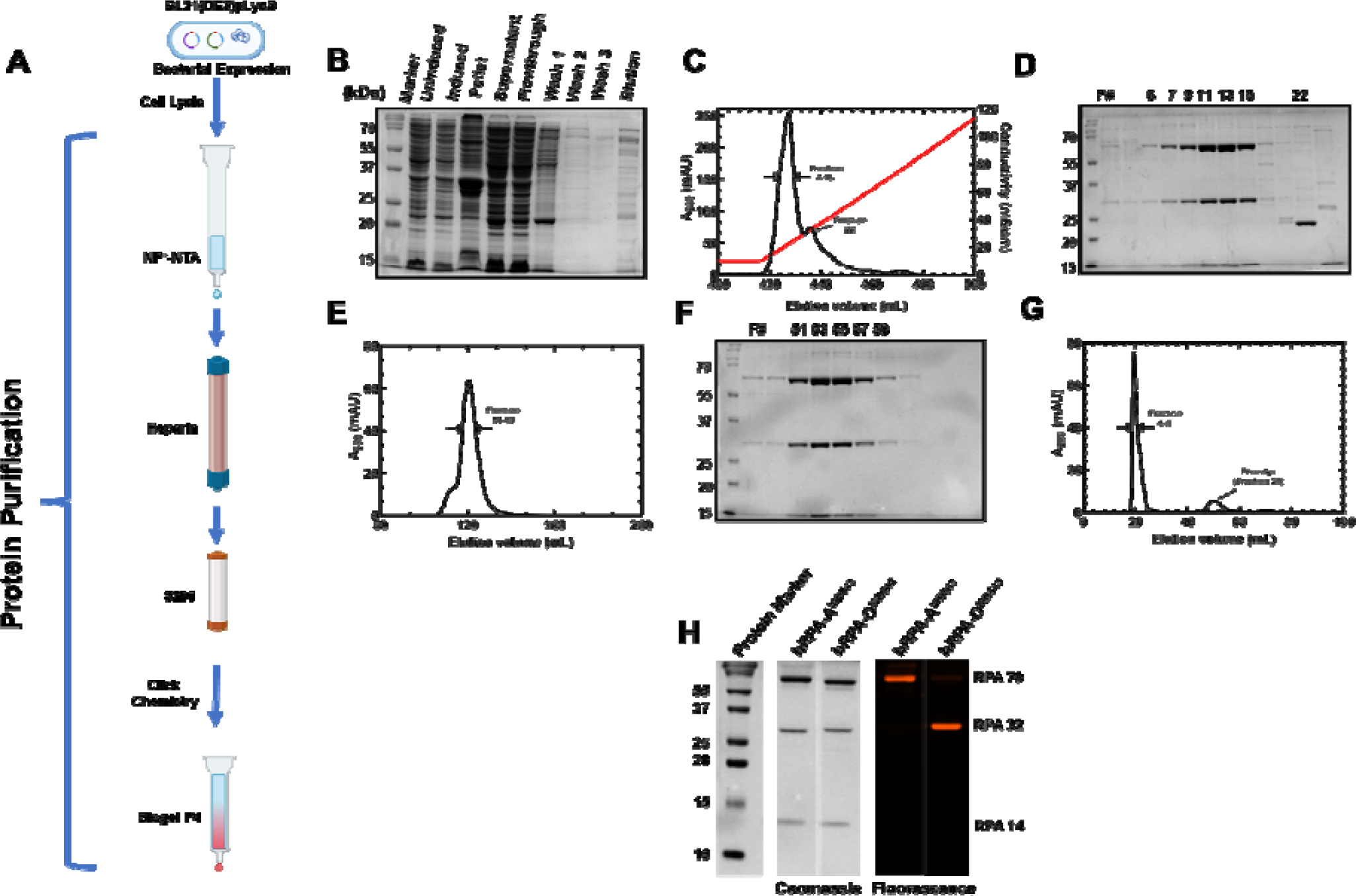
Overproduction, purification, and fluorescent labeling of RPA at DBD-A or DBD-D subunits. (**A**) Workflow to generate the fluorescent versions of RPA. BL21(DE3)pLysS cells were co-transformed with RPA-S215TAG-p11d/RPA-W107TAG-p11d (cDNA sequence) and pDule2-pCNF (t-RNA) plasmids. Transformants were grown and supplemented with 4AZP during induction with IPTG. Recombinant RPA carrying 4AZP (RPA^*4AZP*^) was purified over multiple chromatographic separation steps as denoted. (**B**) SDS-PAGE analysis of fractions during overproduction and the first affinity purification. (**C**) Chromatogram of RPA^*4AZP*^ separation on Heparin and (**D**) SDS-PAGE analysis of fractions eluted from Heparin separation. (**E)** Chromatogram of RPA^*4AZP*^ separation on S200 and (**F**) SDS-PAGE analysis of fractions eluted from size exclusion chromatography. (**G**) Chromatogram of the labeling reaction of RPA^*4AZP*^ and fluorophore separated over a Biogel-P4 column. (**H**) Coomassie and fluorescence images of SDS-PAGE analysis of purified human RPA-DBD-A^MB543^ and RPA-DBD-D^MB543^. All three subunits of RPA are observed in the Coomassie-stained image and only the subunit carrying the fluorophore is visualized in the fluorescence scan.

### Labeling RPA^4AZP^ with click chemistry-based fluorophores

10 μM of RPA^*4AZP*^ (0.5 mL) was mixed with 2X molar excess of the dibenzocyclooctyne-MB543 (DBCO-MB543; Click Chemistry Tools Inc.) dye and incubated at 4 °C for 1.5 h with gentle rocking in dark. Unreacted free dye was separated from the labeled protein using a Biogel-P4 column (40 cm ∼ 2 cm) equilibrated with RPA storage buffer. Eluates carrying fluorescently labeled RPA elutes first and were analyzed on a 12% SDS-PAGE gel and imaged using a iBright 1500 imager (Invitrogen Inc.; **Figure 2G & H**). While all three RPA subunits are observed in the Coomassie stained image, only the subunit carrying the fluorophore is picked up in the fluorescence channel. Concentration and labelling efficiency were calculated using the extinction coefficients of 87,410 M^-1^cm^-1^ and 105,000 M^-1^cm^-1^ for RPA and DBCO-MB543, respectively (**Table 5**). We recommend using the free dye eluting after RPA as a good reference to calculate dye contribution at 280 nm under these buffer conditions. The RPA^*MB543*^ concentration and labeling efficiency were determined as follows:

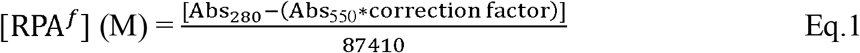

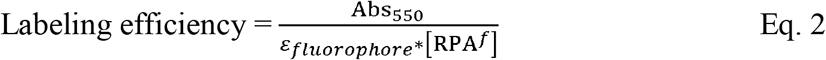

**Table 5.** Extinction coefficient and correction factor for fluorescent dyes:

| Dye | $\epsilon_m$ | Correction Factor |
| --- | --- | --- |
| DBCO-MB543 | 105,000 | 0.08 |
| DBCO-Cy3 | 150,000 | 0.08 |
| DBCO-Cy5 | 250,000 | 0.017 |

The procedure for labeling RPA is similar when using other DBCO-conjugated fluorophores such as Cy3 or Cy5. Please use the extinction coefficients appropriate for the fluorophore used (Cy3 = 150,000 M^-1^cm^-1^ and Cy5 = 250,000 M^-1^cm^-1^; **Table 5**). Fluorescent human RPA is stable for ∼6 months at -80 °C and we do not recommend subjecting the sample to multiple freeze-thaw cycles.

### Measurement of ssDNA binding activity using fluorescence anisotropy

5′-FAM labeled (dT)_35_ ssDNA was diluted to 30 nM in reaction buffer (50 mM Tris-acetate pH 7.5, 50 mM KCl, 5 mM MgCl_2_, 10% v/v glycerol, and 1 mM DTT) and taken in a 10 mm pathlength cuvette (Starna Cells Inc). The temperature was maintained at 23 °C as the fluorescein labelled-ssDNA molecules were excited with vertically polarized 488 nm light and emission was collected using a 520 nm bandpass filter in parallel and perpendicular orientations in PC1 spectrofluorometer (ISS Inc.). The samples were taken in triplicate and five consecutive measurements were averaged for each data point as G-factor corrected anisotropy values were measured after sequential addition of RPA. The titration continued until a steady state was achieved i.e., the anisotropy values plateaued. The raw anisotropy values were corrected for the reduction in quantum yield of the fluorescein moiety upon RPA binding, brought about the proximity based quenching effects of the bound protein molecule as follows:

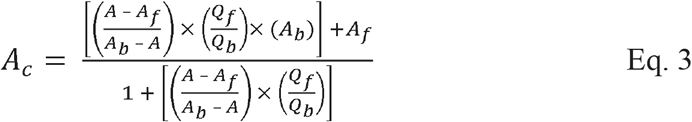

Where, 1) *F*_*b*_, and *F*_*f*_ are the bound, and free concentrations of the FAM-labeled fluorescent ssDNA in nM, 2) *Q*_*b*_, and *Q*_*f*_ are the fluorescence quantum yields of the bound and free form of the FAM-labeled fluorescent ssDNA (arbitrary units), 3) *A*_*b*_, and *A*_*f*_ are the anisotropy values of the bound and free forms of the FAM-labeled fluorescent ssDNA, 4) A, is the measured anisotropy, and 5) *A*_*c*_ is the corrected anisotropy value. Care was taken to limit the maximal dilution to <5% of initial volume and the protein, and ssDNA concentration was corrected for this effect. Due to extremely tight binding between RPA and (dT)_35_ (i.e., also known as stoichiometric binding which is characterized by the lack of free protein during the rising phase of the titration), no attempt was made to fit an equilibrium binding isotherm to the data points.

#### FEncAA assay to capture ssDNA binding induced changes in fluorescence

Another key goal for our approach is that a change in fluorescence (PIFE) must be generated when RPA binds to ssDNA. These PIFE signals will depend on the position of the probe and the relative change in quantum yield of the fluorophore attached. MB453 is an excellent dye to produce PIFE changes and is also relatively photostable. To capture the changes in PIFE, (dT)_35_ was added in a stepwise manner to 100 nM RPA-A^*MB543*^ or RPA-D^*MB543*^ taken in reaction buffer in a 3 mm pathlength cuvette and MB543 fluorescence emission was recorded using PC1 spectrofluorometer. The sample was excited with 530 nm light, and the emitted photons were collected as a function of wavelength in an emission scan. The temperature of the sample was maintained at 23 °C. Following a 2 min incubation after the addition of protein, the fluorescence emission scan was re-acquired. To minimize photobleaching, the excitation shutter was only open during data acquisition. The fluorescence values were corrected for the effects of dilution, photobleaching, and inner filter effects using Eq. 4.

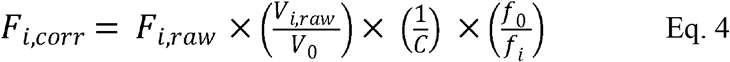

Where, *F*_*i*_,_*raw*_, and *F*_*i*_,_*corr*_ are the measured, and corrected MB543 fluorescence values after each (i^th^) addition of nucleic acid stock. *V*_0_ is the initial volume of the solution before titration, *V*_*i*_ is the volume after after addition of the i^th^ aliquot. *f*_0_, and *f*_*i*_ are the initial fluorescence and fluorescence of RPA due to photobleaching alone under identical solution conditions. C or the correction factor for the inner filter effect was assumed to be 1. The corrected data was plotted as the absolute change in fluorescence normalized to the initial fluorescence 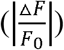 or (fraction enhancement) on Y-axis, with [nucleotides] / [Protein] ratio on the X-axis.

### Stopped-flow experiments to measure Trp quenching and MB543 fluorescence changes

To capture the kinetics of ssDNA binding stopped flow fluorescence experiments were performed on an SX20 instrument (Applied Photophysics Inc.). 100 nM RPA or RPA variants were rapidly mixed with various concentrations of ssDNA (dT)_35_ oligonucleotides (0 – 400 nM) and changes in intrinsic Trp fluorescence was measured by exciting the sample at 290 nm and emission collected using a 350 nm LP filter. Reactions were carried out at 25 °C in reaction buffer (30 mM HEPES pH 7.8, 100 mM KCl, 6% v/v glycerol, 5 mM MgCl_2_, and 1 mM β-mercaptoethanol). Data were fit to a single-exponential plus linear equation to obtain observed rate constants:

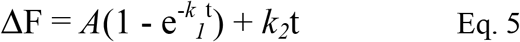

Where F is the Trp fluorescence signal, A is the amplitude of fluorescence change, and *k*_*1*_ and *k*_*2*_ are rates of the two phases. Similar stopped flow experiments were performed to capture the kinetics of RPA^MB543^ variants (RPA-A^MB543^ or RPA-D^MB543^). In this case, 100 nM of the fluorescent RPA from one syringe was mixed with increasing concentrations of (dT)_35_ ssDNA from the other syringe. Changes in MB543 fluorescence were captured by exciting the sample at 535 nm and emission captured using a 555 nm LP filter. Data were fit as above using Eq. 5.

### Stopped-flow FRET assay to monitor the formation of RPA-ssDNA nucleoprotein filaments

A Förster resonance energy transfer (FRET) based experiment was developed to investigate the pre-steady state kinetics of nucleoprotein formation activity of RPA. Here, assembly of multiple fluorescent RPA molecules on a single ssDNA substrate is reported on. RPA molecules are labeled on either DBD-A (Cy5) or DBD-D (Cy3). When these molecules are situated adjacent to each other, an increase in FRET-induced Cy5 fluorescence is observed. To measure the change in Cy5 fluorescence, 200 nM RPA-A^*Cy5*^ and 200 nM RPA-D^*Cy3*^ were rapidly mixed against 90 nM (dT)_97_ in a SX20 stopped flow instrument (Applied Photophysics Inc.). Samples were excited at 535 nM (Cy3 wavelength) and Cy5 emission was captured using a 645 nm long-pass filter.

### Single molecule fluorescence assay to monitor the binding and interactions between two RPA molecules

smFRET-based analysis of RPA binding, diffusion, and mutual interaction on a immobilized (dT)_66_ DNA was carried out by injecting the fluorescently labeled RPA i.e. 150 pM of each RPA-D^*Cy3*^ and RPA-A^*Cy5*^ into an optical cell built over TIRF-microscope; an inverted, objective based total-internal-reflection fluorescence microscope (TIR-FM; IX71 Olympus). TIR excitation was achieved through an oil-immersion objective (100X Olympus UplanApo Numerical Aperture 1.5). The sample was excited with a 532 nm laser, and the emitted fluorescence was split into two channels using Optosplit II (Cairn-Research, UK); Cy3 and Cy5 emissions were collected one half of the chip of the same electron-multiplying charge-coupled device (EM-CCD) camera (iXon Ultra DU-897U-CS0). Images were acquired with 150 ms frame time. Donor and Acceptor images were subpixel registered in Fiji using an image of multicolor beads as fiducial markers. Next, background corrected intensity traces were extracted from immobilized single-molecule spots using Matlab scripts. FRET values were calculated for each fluorescent spot, namely ratio between the acceptor intensity and the sum of the intensities of the donor and acceptor, corrected for crosstalk and cross-excitation. The single molecule FRET v time traces were analyzed in vbFRET to ascribe FRET states and dwell times.

#### Glass coverslips for single-molecule imaging were cleaned and coated as follows

Briefly No. 1.5 thickness Gold Seal coverslips (22 x 60 x 0.17) were sonicated in deionized water, 2% Micro-90, 200-proof ethanol, and finally in 1N KOH all at 60 °C for 15 min. each interspersed with exhaustive washes with deionized water. After drying under a stream of filtered N_2_, the coverslips were first coated with Vectabond (1% v/v Vectabond in 1:18 methanol/acetic acid mixture) followed by washing with deionized water. Next, dried coverslips were incubated overnight with a freshly prepared 18:1 mixture of mPEG-SVA and Biotin-PEG-SVA in 0.1 M NaHCO_3_. Nonspecific binding, which was measured by flowing in non-biotinylated Cy3-or Cy5-labeled DNA or by flowing in biotinylated fluorescent DNA substrate in absence of neutravidin, was <1 %. All measurements were performed at room temperature in buffer (50 mM Tris-acetate pH 7.5, 50 mM KCl, 5 mM MgCl_2_, 10% v/v glycerol, 1 mM DTT, and 0.2-0.5 mg/mL BSA) supplemented with oxygen scavenging solution i.e. 0.1 mg/mL glucose oxidase, 0.2 mg/mL catalase, and 0.4% (wt/wt) β-D-glucose (or 0.1 mM PCD / 10 mM PCA), and 2.5 mM trolox to reduce blinking. To a neutravidin (0.2 mg/mL) coated slide, biotin tagged (dT)_66_ were added at 50-100 pM concentration to arrive at optimal single-molecule number-density of ∼1.2 spot per 3.5 mm^2^ area on the EM-CCD chip.

## Results and discussion

### Generation of fluorescent human RPA carrying non-canonical amino acids

Using the positions that worked well for *S. cerevisiae* RPA as a blueprint, Ser215 in DBD-A (RPA70) and Trp107 in DBD-D (RPA32) are suitable for engineering non-canonical amino acids in human RPA (**Figure 1**). Excellent site-specific incorporation is achieved using 4AZP as the ncAA of choice and copper-free click chemistry is ideal to tether fluorophores onto the desired domain (**Figure 2h**). As with most GCE incorporation strategies, the yields of human RPA carrying 4AZP are lower than for the wildtype protein. Typically, our yields for the RPA^4AZP^ are ∼1.2 mg/L culture. Two important points to note when using our human RPA proteins as probes. The first is that RPA-A^4AZP^ and RPA-D^4AZP^ proteins carry a poly-histidine (6X) tag on the C-terminus of the RPA70 and RPA32 subunits, respectively. We have not attempted to engineer a cleavable affinity tag as the DNA binding activity was unperturbed (**Figures 3 & 4**). Second, the labeling efficiencies for RPA-A^4AZP^ and RPA-D^4AZP^ are ∼55% and ∼35%, respectively. These are largely defined by the local accessibility of the 4AZP group on the protein and the limitations of the click chemistry DBCO groups used. A balance between time of reaction versus protein stability needs to be considered and conditions described here have been extensively optimized for human RPA. If this approach is to be applied for other proteins of interest, careful optimization of reaction conditions is recommended.

**Figure 3.**
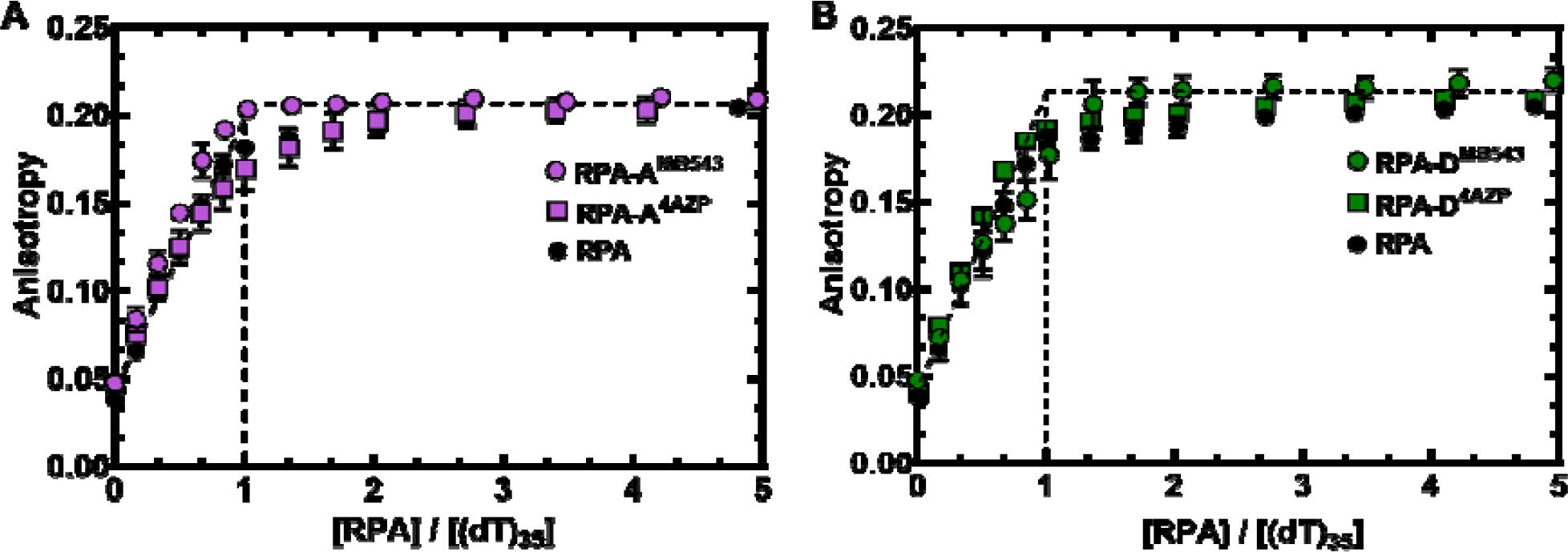
RPA^*4AZP*^ and RPA^*MB543*^ bind to ssDNA with affinity comparable to RPA. Change in fluorescence anisotropy of a 5_′_-FAM-labelled (dT)_35_ ssDNA substrate was monitored as a function of DNA concentration to compare the binding stoichiometry of (**A**) RPA, RPA-A^*4AZP*^, and RPA-A^*MB543*^ and (**B**) RPA, RPA-D^*4AZP*^ and RPA-D^*MB543*^. All versions of RPA bind to ssDNA with high affinity and comparable stoichiometry.

### Fluorescent human RPA show no loss in ssDNA binding properties

One of our first objectives was to produce a version of fluorescent human RPA that retains ssDNA activity similar to (or as close to as feasible) the wildtype protein. In steady state ssDNA binding anisotropy experiments, all versions of human RPA (RPA, RPA-A^4AZP^, RPA-D^4AZP^, RPA-A^MB543^, and RPA-D^MB543^) show stoichiometric ssDNA binding activity (**Figure 3**). This is hardly surprising since there are four high affinity DNA binding domains in RPA and collectively, they bind to ssDNA with very high affinity. A better measure of the ssDNA binding properties is a kinetic assessment of the overall change in intrinsic tryptophan (Trp) fluorescence. We monitored the change in Trp fluorescence using a stopped flow assay and obtained the binding and dissociation rates for RPA to a (dT)_35_ ssDNA substrate (**Figure 4**). Human RPA and the two fluorescent versions bound to ssDNA with similar rates and display comparable k_on_. These findings satisfy one of the key criteria for these fluorescent RPA probes: there is no measurable loss in DNA binding properties.

**Figure 4.**
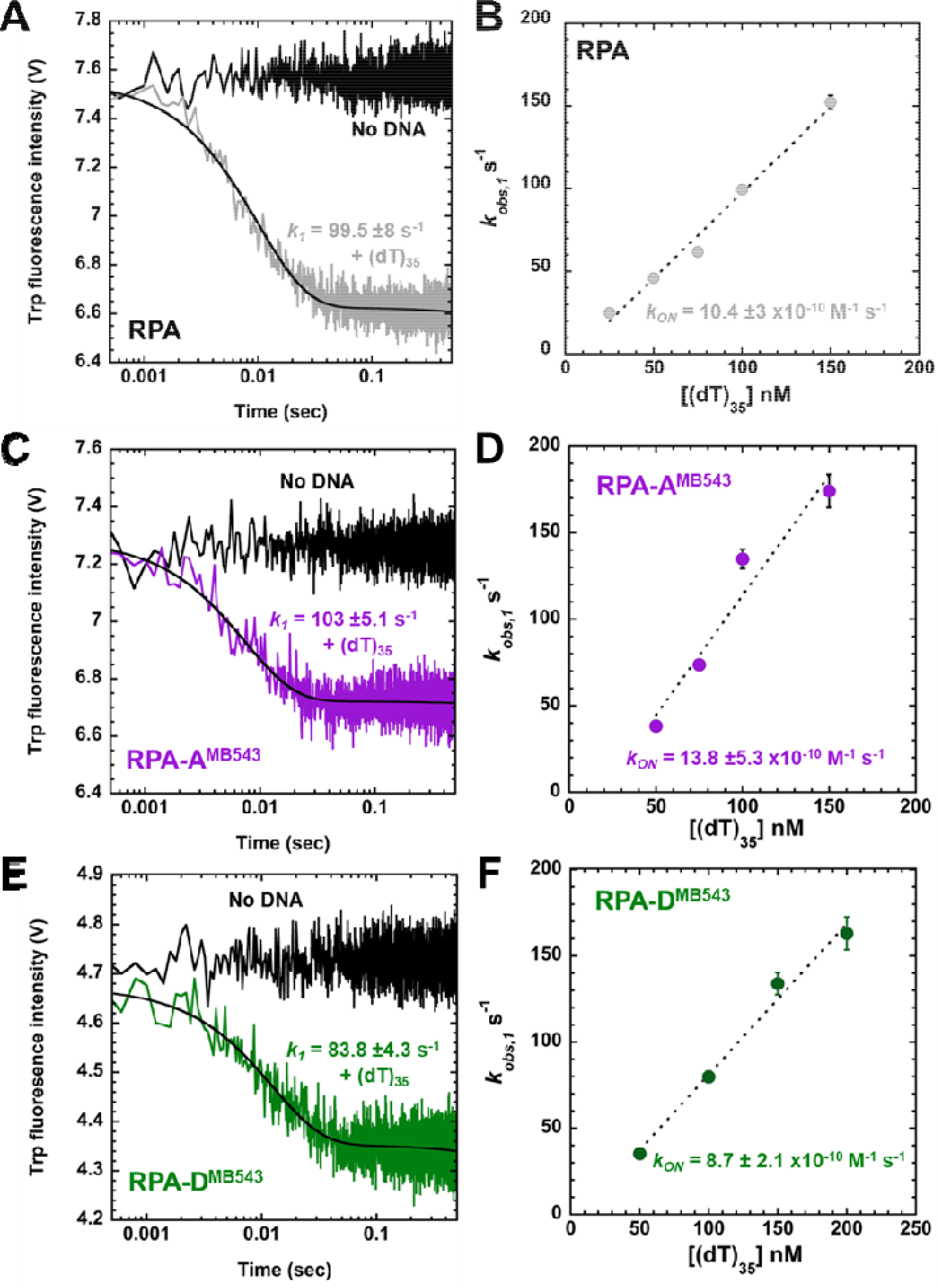
Kinetics of ssDNA binding reveal the human RPA fluorescent proteins to be proficient for DNA interactions. Stopped flow experiments were performed by mixing (**A, B**) RPA, (**C**,**D**) RPA-A^MB543^, or (**E**,**F**) RPA-D^MB543^ with increasing concentrations of (dT)_35_ ssDNA and the change in intrinsic Trp fluorescence was captured. Plots of the k_obs_ versus DNA concentration yield k_ON_ parameters that are comparable for all three RPA proteins.

### ssDNA binding produces an enhancement in fluorescence

Another unique feature of our probe design is to produce a change in fluorescence when human RPA interacts with ssDNA. ssDNA binding leads to a change in the local microenvironment of the domains. MB543 is an environmentally sensitive fluorophore whose quantum yield is influenced by the local dipole moment and micro-viscosity (among other factors). To capture changes in fluorescence, RPA-A^*MB543*^ or RPA-D^*MB543*^ was titrated against (dT)_35_ ssDNA and the fluorescence quantum yield was plotted against the ssDNA concentration. Both RPA-A^*MB543*^ and RPA-D^*MB543*^ produce observable changes in MB543 fluorescence upon binding ssDNA. The amplitude of change is much larger for RPA-A^*MB543*^ (35%) compared to RPA-D^*MB543*^ (8%) owing to the position of the fluorophores on the respective DBDs and also due to differential changes in the local environment of the respective domains (**Figure 5A & D**). In the ssDNA titrations, both fluorescence signal changes report on stoichiometric ssDNA interactions for RPA-A^*MB543*^ and RPA-D^*MB543*^ (**Figure 5B & D**).

**Figure 5:**
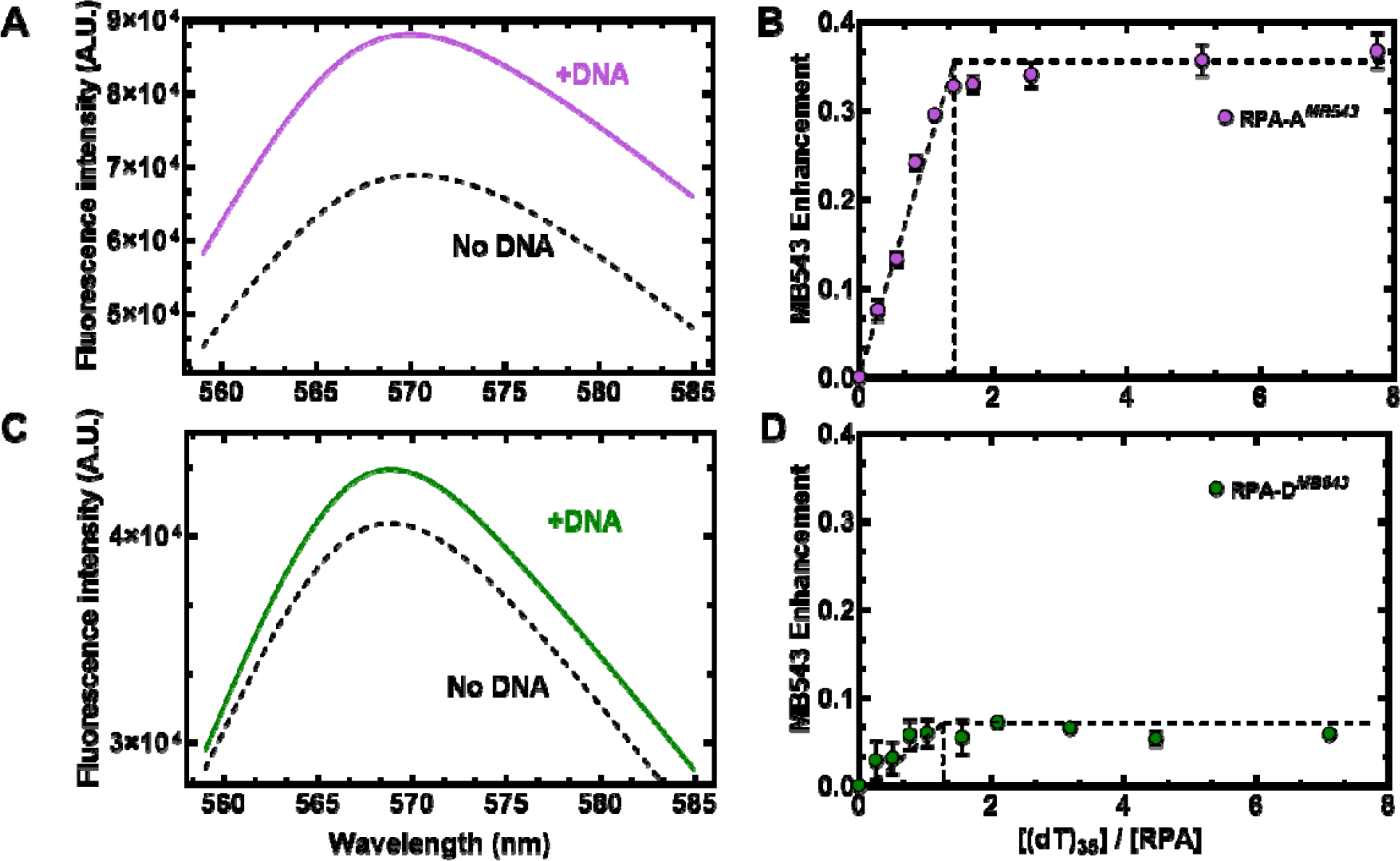
FEncAA assay to quantitatively monitor the binding of (dT)_35_ ssDNA to RPA-A^*MB543*^ or RPA-D^*MB543*^. Fluorescence spectra showing the quantum yield of the environmentally sensitive fluorophore MB543 positioned in (**A**, **B**) RPA-A^MB543^ versus (**C**, **D**) RPA-D^MB543^ in the absence and presence of (dT)_35_ ssDNA. Quantitation of the changes in fluorescence shows stoichiometric ssDNA binding by both versions of RPA. The amplitude of fluorescence change is greater for RPA-A^MB543^ compared to RPA-D^MB543^ suggesting more solvent exposure for the fluorophore on RPA-A^MB543^.

### Human RPA probes report on the dynamics of individual DNA binding domains

One objective in designing these site-specific FEncAA probes is to experimentally quantititate the DNA binding dynamics of a single DBD within the context of full length RPA. As a proof of concept, we performed stopped flow experiments where the fluorescence of RPA-A^*MB543*^ or RPA-D^*MB543*^ was tracked when rapidly mixed with (dT)_35_. For RPA-A^*MB543*^ two binding phases are observed (**Figure 6A**). Similarly, for RPA-D^*MB543*^ two binding phases are observed (**Figure 6B**). The observed rates are roughly similar, although we caution that the signal to noise is significantly higher for RPA-D^*MB543*^ and thus we refrain from making mechanistic conclusions at this juncture. Nevertheless, both DBDs appear to have a rapid initial engagement step followed by a slower remodeling step. There are many applications for these human RPA probes and the ability to site-specifically position the fluorophores opens a toolkit of ensemble and single-molecule experiments (described below).

**Figure 6.**
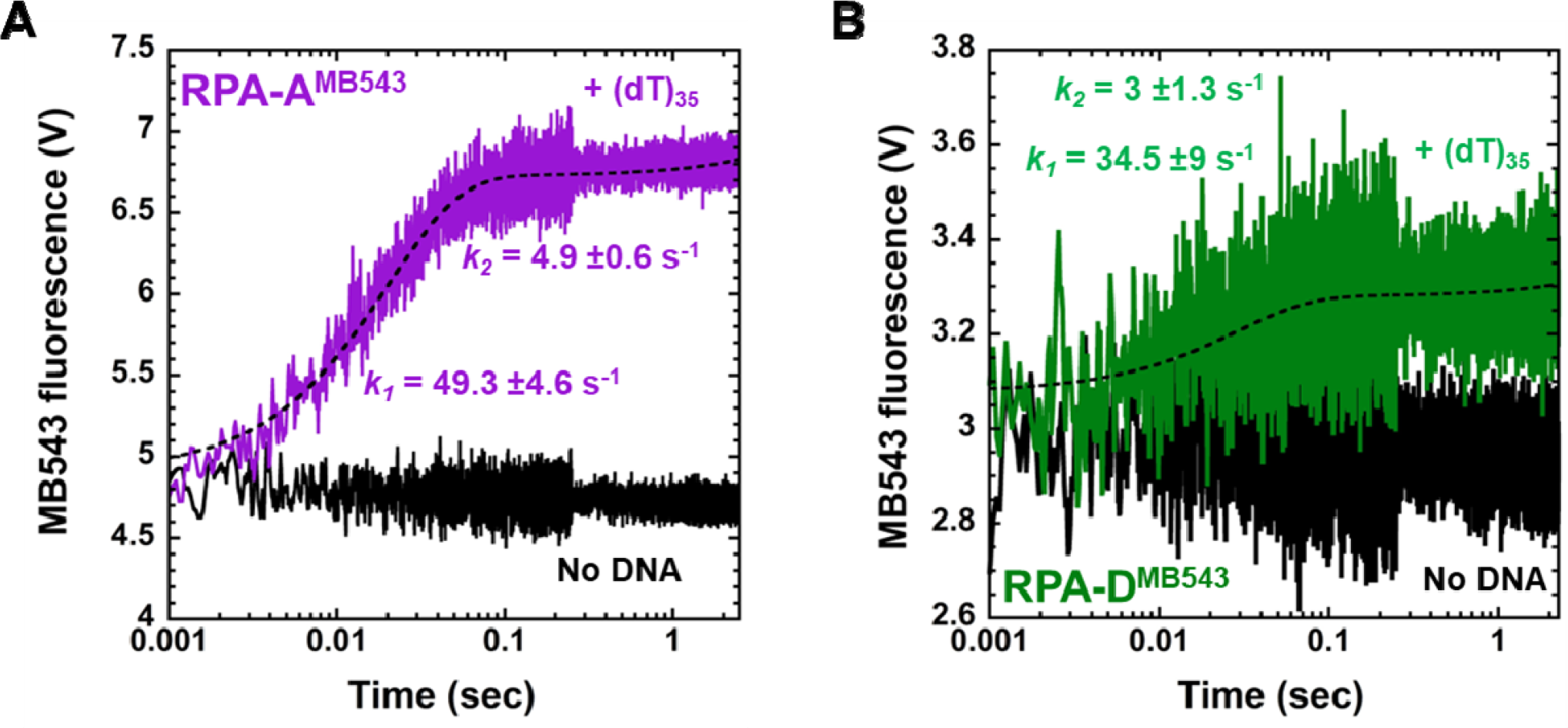
Site-specific fluorophores report on ssDNA binding dynamics of select DNA binding domains. Stopped flow analysis of (dT)_35_ binding to (**A**) human RPA-A^MB543^ or (**B**) RPA-D^MB543^ show bi-phasic changes in fluorescence. The data show that the two domains possess a rapid DNA binding phase followed by a slower remodeling phase. Comparison of the observed rates reveal a marginal faster binding of DBD-A.

### Kinetics of RPA-ssDNA nucleoprotein filament formation

On longer ssDNA, multiple RPA molecules assemble to form nucleoprotein filaments to reduce formation of secondary structures. Using our fluorescent human RPA probes we can assess how RPA assembles into such larger complexes. We assayed the kinetics of RPA binding to a longer piece of (dT)_97_ ssDNA using a stopped flow. Given the site-size of ∼20 nt/RPA, about 4-5 RPA molecules can be bound this substrate. For this experiment, we generated Cy5 or Cy3 labeled human RPA. Human RPA-D^Cy3^ and RPA-A^Cy5^ were generated and equimolar amounts of both these fluorescent RPA were rapidly mixed with (dT)_97_ ssDNA. Since RPA binds with defined 5′->3′ polarity^8,39,40^, with DBD-A aligned towards to the 5′ side (**Figure 7A**), Cy3 on DBD-D from one RPA molecule will be positioned adjacent to DBD-A from a neighboring RPA molecule. Such an arrangement would produce an increase in a FRET when Cy3 is excited and change in Cy5 fluorescence is monitored. An example of this experiment is shown (**Figure 7B**). We have used such FRET experiments to monitor assembly and disassembly of RPA on longer DNA substrates.^41,42^ Such experiments would be powerful to investigate the role of post-translational modifications or mutations in RPA and to assess the role of RPA-interacting proteins in remodeling RPA.

**Figure 7.**
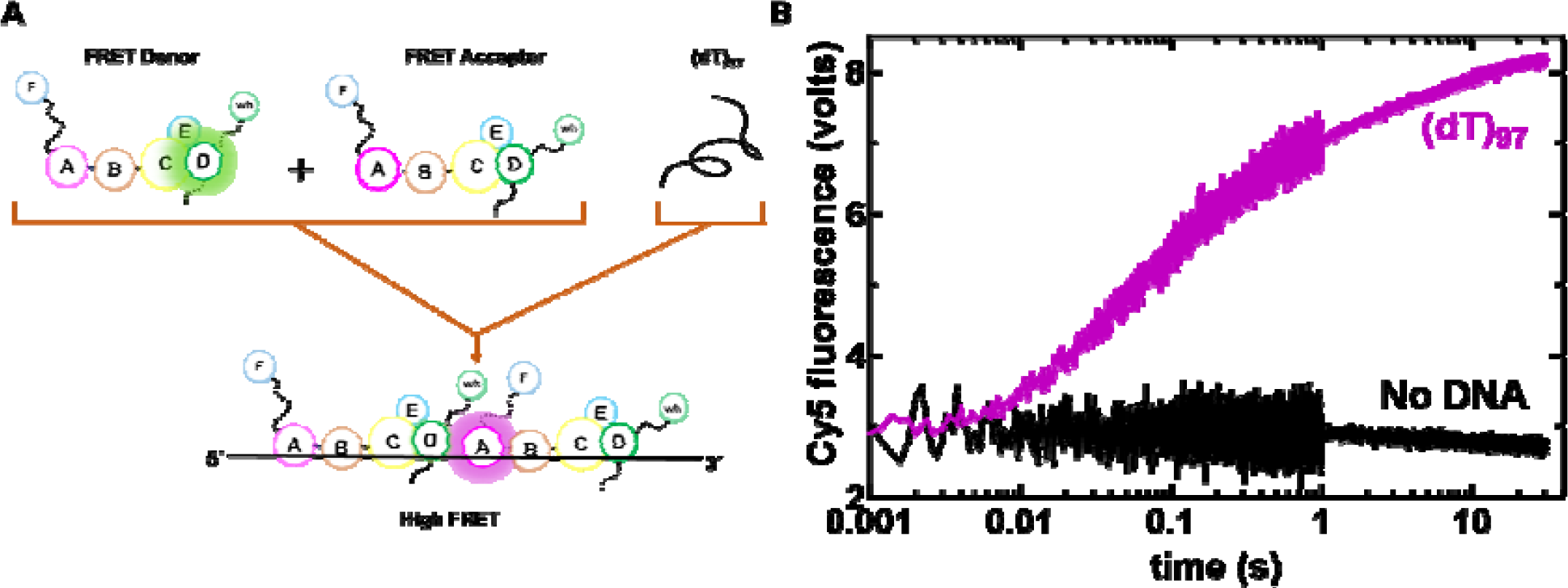
Pre-steady state kinetics of RPA-ssDNA filament formation. (**A**) Cartoon depicting assembly of RPA-(dT)_97_ nucleoprotein filaments. The signal change arises from energy-transfer between adjacently bound RPA-D^*Cy3*^ and RPA-A^*Cy5*^ molecules. (**B**) Donor is excited at 530 nm and energy transfer (FRET) between Cy3 and Cy5 can be observed at 678 nm (magenta). Change in Cy5 fluorescence is observed in the presence of ssDNA. This experiment can be utilized to capture the kinetics of RPA filament formation.

### Single molecule assay to monitor binding & transient interactions between two RPA molecules

When two or more molecules of RPA bind to a sufficiently long piece of ssDNA, it results in the formation of a nucleoprotein filament as described above. It’s not clear if adjacent RPA molecules bind or interact with each other as a part of nucleoprotein filament. However, there have been suggestions for yeast RPA acting with weak cooperatively in binding to ssDNA^28^. Here we present a single-molecule total internal reflection microscopy (smTIRM) assay to directly measure the lifetime of interactions of two RPA molecules as a part of nucleoprotein filament. Two molecules of human RPA labeled with either Cy3 on DBD-D or Cy5 on DBD-A are flown onto a biotin-(dT)_66_ substate anchored to the surface of a quartz slide (**Figure 8A**). Fluorescence signals are captured for the donor (Cy3) and acceptor (Cy5) and FRET changes are calculated from individual single-molecule donor and acceptor trajectories (**Figure 8B**). Data shows anti-correlated intensity fluctuations and clearly depict high FRET events which last for a few seconds (blue trace in **Figure 8B**). The lifetime (δt) of these high FRET events reflects interaction time between two adjacent RPA molecules as they laterally diffuse on the ssDNA lattice and collide. The overall FRET histogram shows that RPA molecules tend to spend most of the time together (<3 nm) in a state of high FRET (**Figure 8C**). In addition to the single-molecule experiments shown above, these fluorescent human RPA probes can be used in C-trap experiments to track RPA dynamics on kilobases long ssDNA substrates. We have used such assays to monitor RPA disassembly by the HELB helicase.^43^

**Figure 8.**
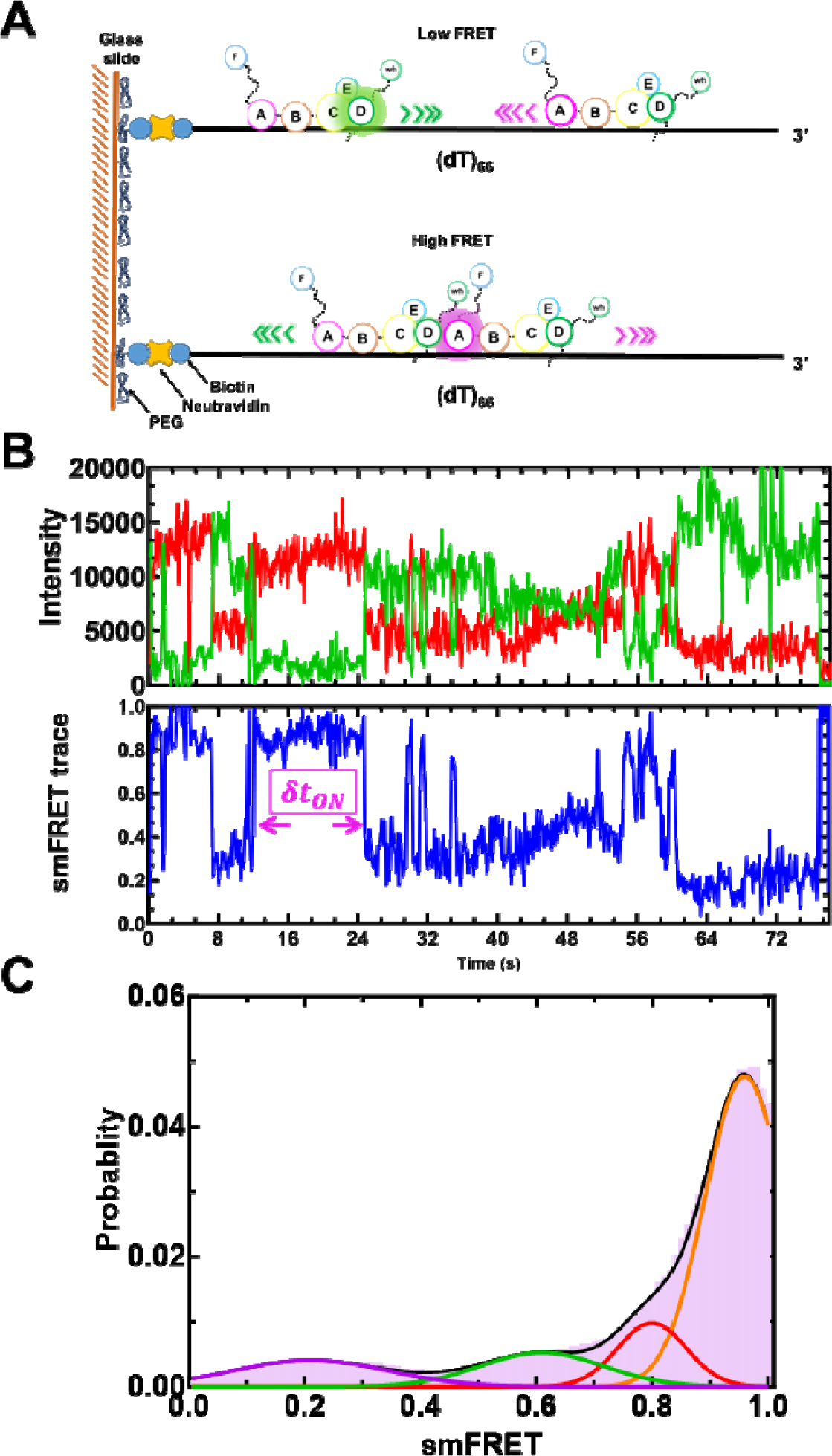
Single molecule assay to estimate the interaction time between two RPA molecules bound to (dT)_66_. (**A**) Cartoon depicts the assembly and dynamics of two RPA molecules labeled (with Cy3 or Cy5 at D or A subunits, respectively) on a surface tethered (dT)_66_ ssDNA molecule. As bound RPA molecules diffuse along the length of DNA collision of two RPA molecules will produce a transient high FRET signal (magenta). The lifetime of these fleeting binding events can be calculated from smFRET trajectories (δt_ON_ ; **B**). (**C**) A smFRET frequency histogram shows relative locations of RPA on (dT)_66_ ssDNA. High FRET indicates that the RPA molecules spend relatively large amount of time close to each other on (dT)_66_ ssDNA under the conditions tested.

## Acknowledgements

This work was supported by grants from the NIH, NIGMS R35 GM149320 to E.A. and R01 GM134081 to C.K.A. S.K., was supported by a fellowship from the NIH, NCI F99 CA274696. AUC experiments to characterize the oligomeric state of RPA was supported by a grant from the NIH Office of the Director S10 OD030343 to E.A.

